## Supplemental figures and tables for "Envelope reconstruction of speech and music highlights unique tracking of speech at low frequencies"

Supplemental information

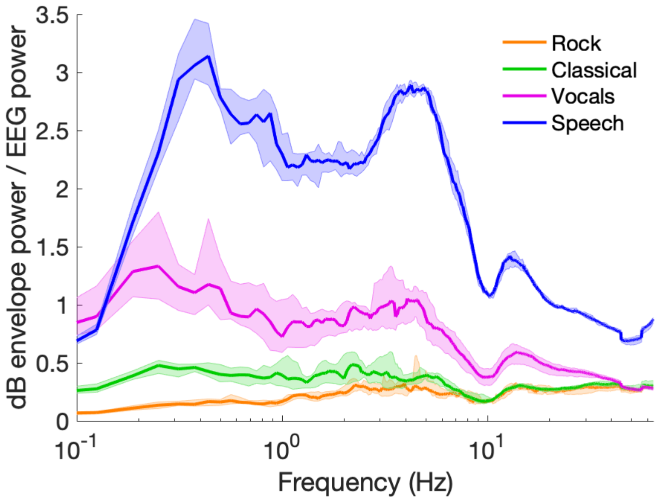


Figure S1: We were interested in examining the relative power spectra of the envelopes, in order to quantify the hypothesis that EEG tracking of the envelope is a direct replicate of the envelope itself, and that the differences in reconstruction accuracy are due to differences in envelope variance at each frequency range. To do this, each dB envelope was zero-centered, and a 16 s moving average was subtracted (equal to the maximum model delay, see "Quantifying cross-frequency model performance" in the Methods). All envelopes were then normalized by the square root of the average variance across all stimuli. Then, the power spectra of individual envelopes were computed. To compute "EEG noise", the EEG data in each trial was averaged across channels, the 16 s moving average was subtracted, and the averaged EEG was z-scored. Then, the power spectrum was computed on the z-scored EEG data. Power spectra of the envelopes were averaged across trials (including all stimulus types) and subjects, and the ratio of the envelope power to EEG noise power was computed, where the denominator of the ratio was using the trial- and subject-averaged EEG spectrum. Lines designate the medians of each stimulus type, and shaded regions designate the 95% quantile of bootstrapped medians.

The Envelope power to EEG power ratio is theoretically proportional to the signal-to-noise ratio of the neural response to the envelope, assuming that the neural response is a scaled version of the dB envelope (the "null hypothesis") and the EEG noise is proportional to the average EEG spectrum. Note that while we have no knowledge of the linear relationship between dB envelope and the neural response, a change in the scaling would multiply these ratios for each stimulus type identically and maintain their relationships to each other. For the purpose of identifying the differences in neural tracking due to envelope variance in each frequency range, one should examine the relative magnitudes of the Envelope power to EEG power between stimulus types, and not the absolute magnitudes.

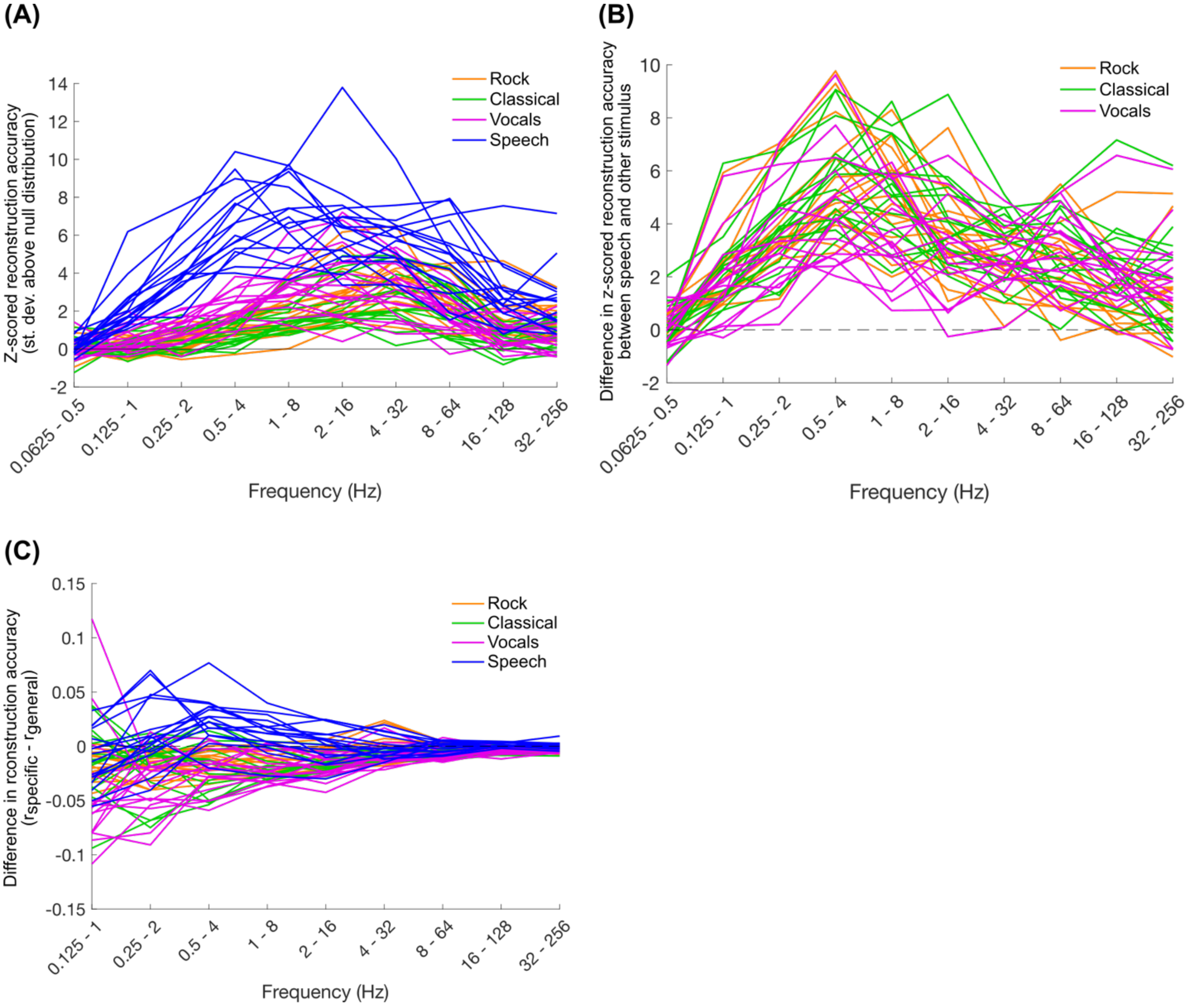


Figure S2: Reconstruction accuracy curves for individual subjects. (A) Z-scored reconstruction accuracies for each stimulus type (compare to Figure 3d). (B) Difference between z-scored reconstruction accuracy for speech and each of the other stimulus types (compare to Figure 3e). (C) Difference between stimulus-specific and stimulus-general reconstruction accuracy (compare to Figure 5a).

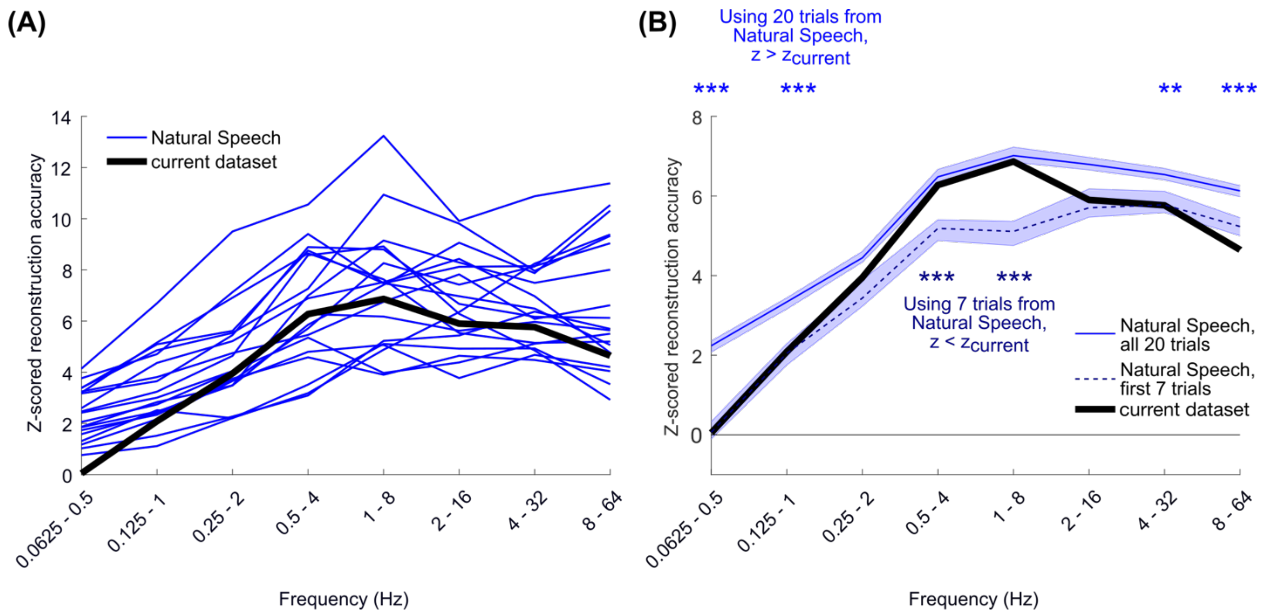


Figure S3: We repeated the frequency-constrained reconstruction accuracy analysis on the Natural Speech dataset (Broderick et al, 2018), in order to validate that the high low-frequency reconstruction accuracies we observed for speech were not specific to the current dataset (compare to Figure 3d). Note that the Natural Speech dataset contained 20 trials of the audiobook, whereas the current dataset in the study only contained the first 6-7 trials (7 for most subjects, see Table S1). (A) Shown are the reconstruction accuracies for all 19 subjects in the Natural Speech dataset, averaged across 20 trials. (B) We looked at reconstruction accuracies using all 20 trials of Natural Speech (blue, same results as A) and only the first seven trials (darker blue, dashed line in B). Wilcoxon's rank-sum test with Bonferroni correction for 16 comparisons was used to compare reconstruction accuracies between datasets; blue shows the comparisons with all 20 trials of Natural Speech, and darker blue shows comparisons is using just the first seven trials (** $p < 0.01$; *** $p < 0.001$). In both instances, reconstruction accuracies were comparable to the current dataset and higher than the reconstruction accuracies for the other stimuli (see Figure 3d, S1). Note, however, that using all 20 trials produces above-chance reconstruction accuracies for the lowest frequency model, 0.0625 – 0.5 Hz. The reconstruction accuracies drop to chance when only seven trials are used. This indicates that the chance performance we observed in the current dataset may not be due to a low-frequency limitation on neural tracking of the speech envelope and may instead be a result of the limited amount of data in this study.

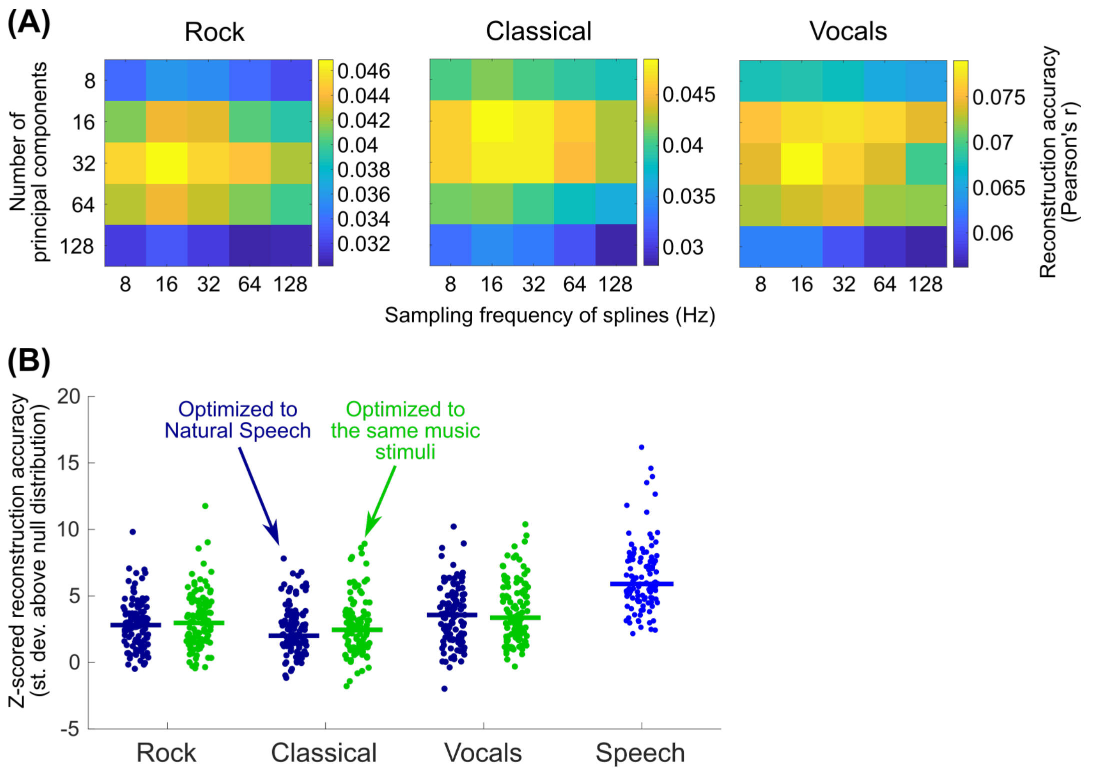


Figure S4: We optimized the hyperparameters of the PCA & spline model (specifically, the sampling frequency of the spline knots and the number of principal components) to a separate speech dataset, and we found that speech envelope reconstruction was better than music for all of the frequency ranges we examined (see Figure 3). Here, we tested if music envelope reconstruction performs as well as speech if we optimize the hyperparameters for the music stimuli. (A) Using the same 500 ms model window as before, we found the optimal hyperparameter pairs for each stimulus type that maximized the average envelope reconstruction accuracy across subjects. These optimal hyperparameters were different than those found for Natural Speech (Rock = 16 Hz spline knots, 32 principal components (PCs); Classical = 16 Hz, 16 PCs; Vocals = 16 Hz, 32 PCs; Natural Speech = 32 Hz, 64 PCs). (B) We then computed the z-scored reconstruction accuracies (as in Figure 3) using these optimal hyperparameters. For each music stimulus, the dark blue dots on the left are the trial-by-trial reconstruction accuracies for all subjects using the Natural Speech hyperparameters (the same datapoints as those used to create Figure 3d), and the green dots on the right are using the music-optimized hyperparameters. The blue dots for the speech z-scored accuracies are based on the Natural Speech hyperparameters. Lines indicate the median values across trials and subjects. Even after optimizing the hyperparameters to the music stimuli, speech envelope reconstruction still outperforms music (Wilcoxon rank-sum relative to speech: $z_{rock} = 9.17$, $z_{classical} = 9.84$, $z_{vocals} = 7.16$, $p < 0.001$ for all comparisons).

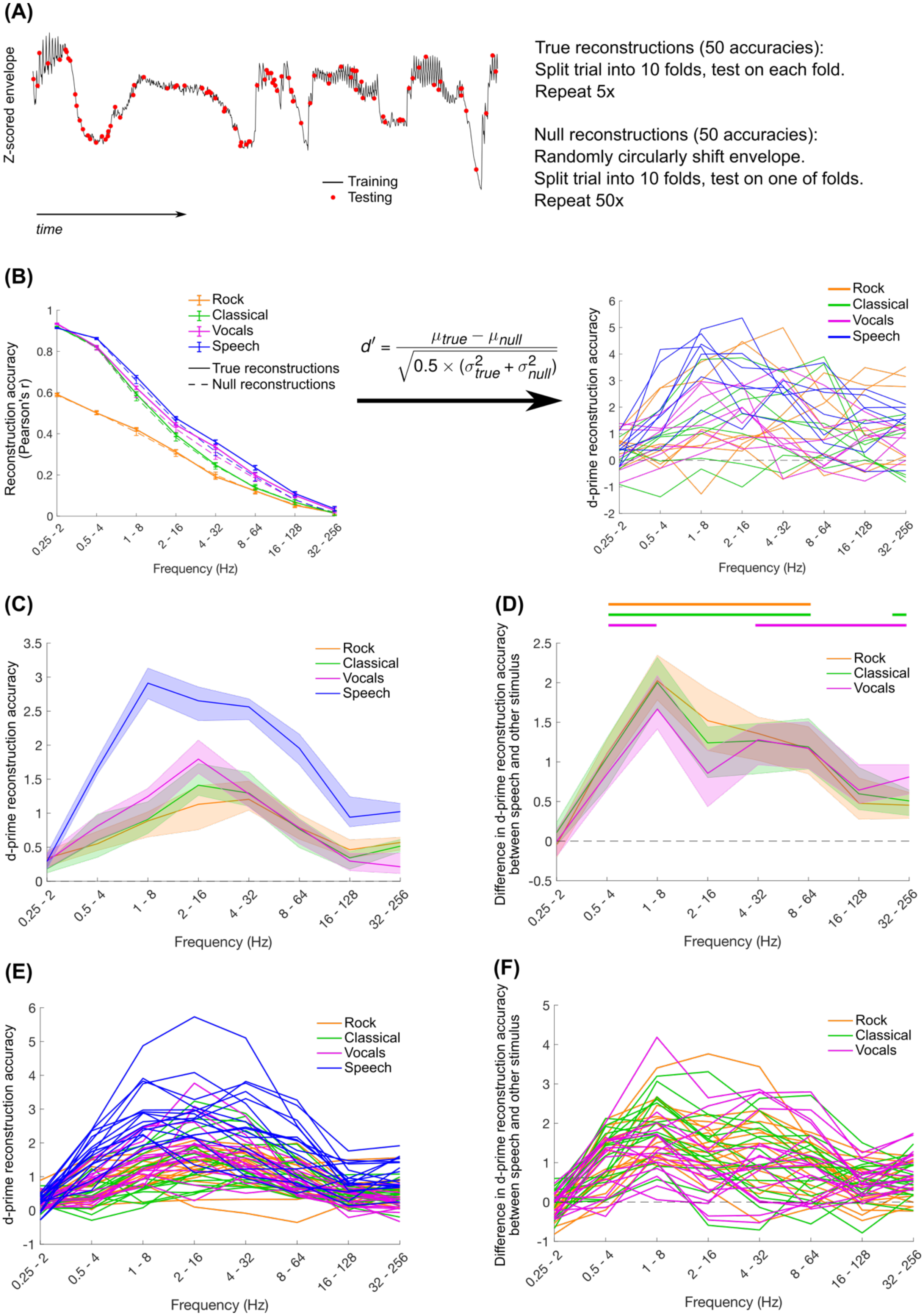

(A)
Z-scored envelope
Training
Testing
time
True reconstructions (50 accuracies):
Split trial into 10 folds, test on each fold.
Repeat 5x
Null reconstructions (50 accuracies):
Randomly circularly shift envelope.
Split trial into 10 folds, test on one of folds.
Repeat 50x
(B)
Rock
Classical
Vocals
Speech
True reconstructions
Null reconstructions
Reconstruction accuracy (Pearson's r)
$d' = \frac{\mu_{true} - \mu_{null}}{\sqrt{0.5 \times (\sigma^2_{true} + \sigma^2_{null})}}$
d-prime reconstruction accuracy
Frequency (Hz)
0.25 - 2
0.5 - 4
1 - 8
2 - 16
4 - 32
8 - 64
16 - 128
32 - 256
(C)
d-prime reconstruction accuracy
Rock
Classical
Vocals
Speech
Frequency (Hz)
(D)
Difference in d-prime reconstruction accuracy between speech and other stimulus
Rock
Classical
Vocals
Frequency (Hz)
(E)
d-prime reconstruction accuracy
Rock
Classical
Vocals
Speech
Frequency (Hz)
(F)
Difference in d-prime reconstruction accuracy between speech and other stimulus
Rock
Classical
Vocals
Frequency (Hz)

Figure S5: We observed higher reconstruction accuracies for speech than the music stimuli (Figure 3d, e) but this could have been due to the higher cross-trial variability for the music stimuli than for the speech stimuli. To control for this, we looked instead at within-trial reconstruction accuracy. (A) To get reconstruction accuracies for each trial, we split the trial into 10 evenly-sized folds, where each fold contained a random sampling of the data in the trial. This was done in order to maximize the consistency in the EEG covariance and envelope spectrum across folds. Then models were fit on all trials with one fold left out and tested on the left-out fold. This was repeated 5 times using a new random sampling of folds each time, giving a total of 50 reconstruction accuracies (Pearson's r) for each trial. To get a null distribution of accuracies, the stimulus envelope was randomly circularly shifted, $1/10^{th}$ of the data was randomly sampled for testing, and the rest of the data was used for training. This was repeated 50 times to get 50 null reconstruction accuracies. (B) Because testing data is highly correlated with training data using this method, both the true and null reconstruction accuracies increase as lower frequencies are used for modeling. To correct this, we computed a d-prime reconstruction accuracy based on the distribution of true and null reconstruction accuracies. (C, D) Firstly, d-prime reconstruction accuracies dropped to zero for the 0.25-2 Hz model. This is a consequence of the reduced amount of data available in each trial; using lower-frequency models (with larger model windows) generated warnings in Matlab indicative of overfitting. But that aside, across all frequency ranges, d-prime reconstruction accuracy was significantly larger than all other music stimuli. Thick lines in D show significance of a permutation test comparing speech d-prime to each of the different stimulus types, $p < 0.001$ with Bonferroni correction for 24 comparisons. Plots (E and F) show the same results as C and D, respectively, for individual subjects. Overall, this indicates that, even when doing within-trial reconstructions to avoid the effects of cross-trial variance, speech is reconstructed better than the music stimuli.

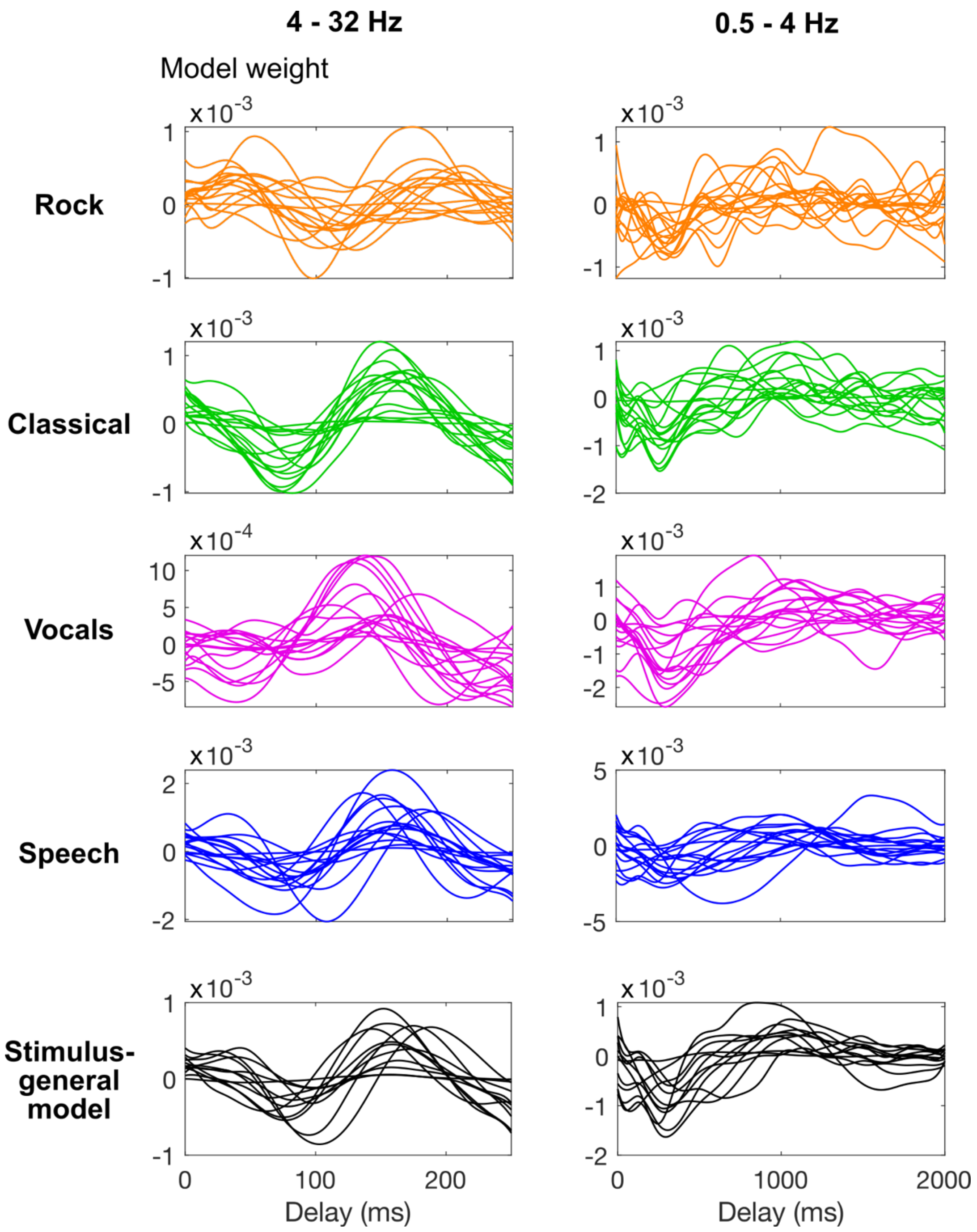


Figure S6: Temporal weights for reconstruction models for each individual subject, averaged across trials, after transforming to a "forward" model (Haufe et al, 2014). The models were converted from basis splines to delays, and then from principal components to EEG channels. The weights shown here were averaged across all 128 EEG channels (compare to Figure 4 and Figure 5b).

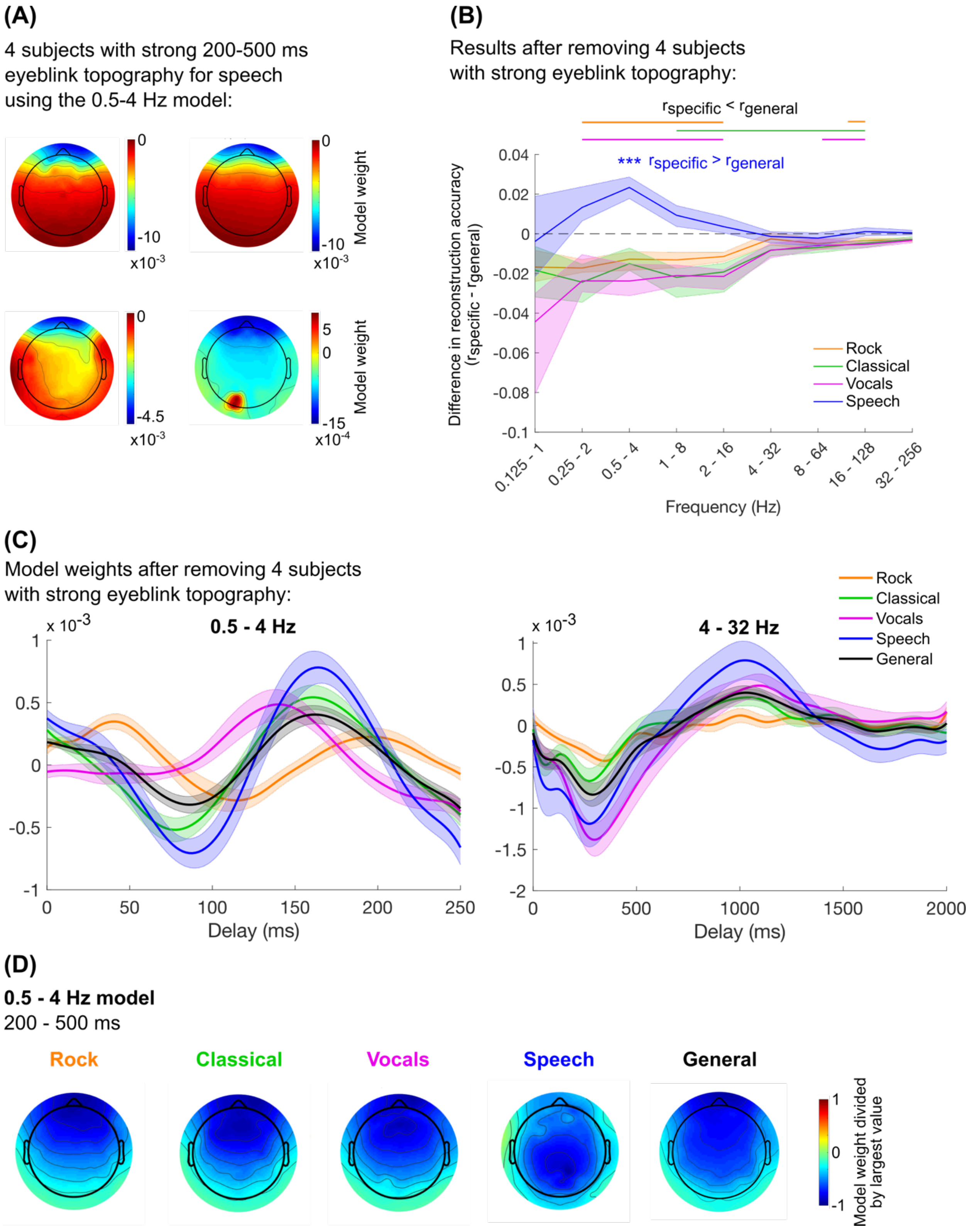

(A)
4 subjects with strong 200-500 ms eyeblink topography for speech using the 0.5-4 Hz model:
0
-10
x10-3
0
-10
x10-3
Model weight
0
-4.5
x10-3
5
0
-15
x10-4
Model weight
(B)
Results after removing 4 subjects with strong eyeblink topography:
rspecific < rgeneral
*** rspecific > rgeneral
Difference in reconstruction accuracy (rspecific - rgeneral)
0.04
0.02
0
-0.02
-0.04
-0.06
-0.08
-0.1
0.125 - 1
0.25 - 2
0.5 - 4
1 - 8
2 - 16
4 - 32
8 - 64
16 - 128
32 - 256
Frequency (Hz)
Rock
Classical
Vocals
Speech
(C)
Model weights after removing 4 subjects with strong eyeblink topography:
x 10-3
0.5 - 4 Hz
1
0.5
0
-0.5
-1
0
50
100
150
200
250
Delay (ms)
x 10-3
4 - 32 Hz
1
0.5
0
-0.5
-1
-1.5
-2
0
500
1000
1500
2000
Delay (ms)
Rock
Classical
Vocals
Speech
General
(D)
0.5 - 4 Hz model
200 - 500 ms
Rock
Classical
Vocals
Speech
General
1
0
-1
Model weight divided by largest value

Figure S7: Based on the frontal topography of the weights that we observed for the 0.5 – 4 Hz model (Figure 4d, 5c), we were concerned that the greater reconstruction accuracies for the speech-specific model compared the stimulus-general model (Figure 5a) might be related to eyeblink artifacts, which were especially prominent in some subjects (for example, if subjects unconsciously timed eyeblinks to envelope onsets in the stimulus). While, to our knowledge, no eyeblink-based speech envelope reconstruction has been reported in the past, a 300-400 ms frontal negativity is indicative of eyeblink contamination in evoked response analyses (Talsma & Woldorff, 2005). (A) We examined the topography of the weights between 200 – 500 ms for each individual subject and found four subjects with topographies strongly indicative of eyeblinks. (B) After removing these subjects from analysis, however, the stimulus-specific model for speech still outperformed the stimulus general model for 0.5-4 Hz (Wilcoxon signed-rank test with Bonferroni correction for 32 comparisons, $p < 0.001$), but this was no longer true for the 1-8 Hz model. (C and D) Additionally, the time course and topographies of the model weights were very similar to what was observed in our analysis using all 16 subjects (compare to Figure 4 and Figure 5).

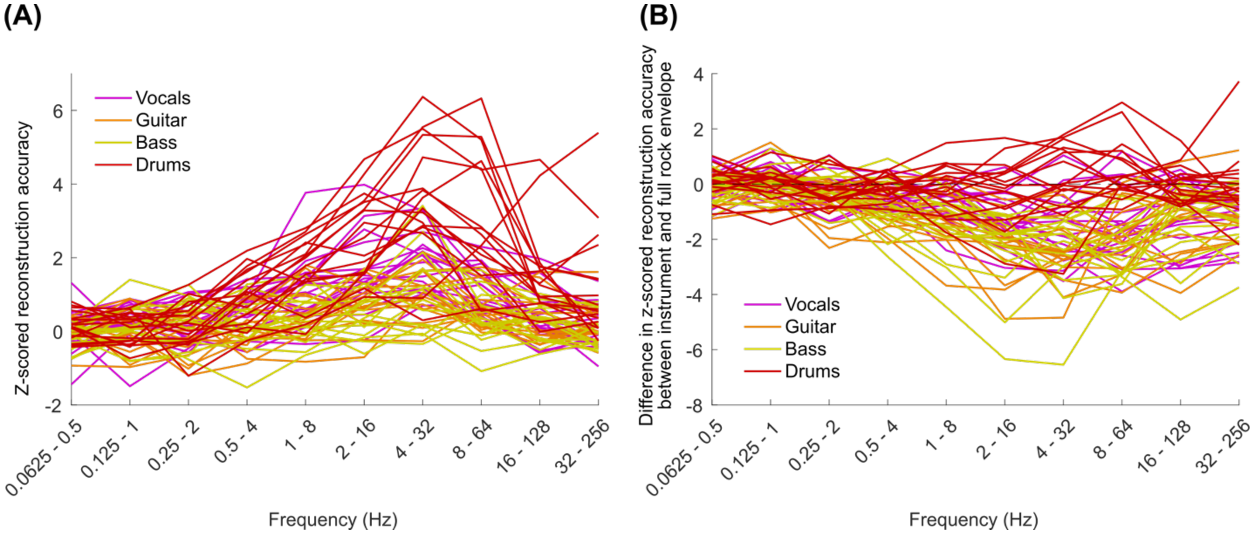


Figure S8: Rock instrument reconstruction accuracies for individual subjects. (A) and (B) are plotted identically to Figure 6a and b respectively.

Table S1: List of stimuli used in this experiment, including stimulus duration and the number of times it was presented across all 16 subjects.

| Title | Artist / Composer | Duration (s) | Number of presentations across all subjects |
|---|---|---|---|
| **Rock** | | | |
| Mr. Brightside | The Killers | 225.68 | 16 |
| Call Me | Blondie | 222.16 | 10 |
| Kids in America | The Muffs | 216.70 | 10 |
| Long Road to Ruin | Foo Fighters | 216.17 | 12 |
| Livin’ On A Prayer | Bon Jovi | 283.08 | 12 |
| Go Your Own Way | Fleetwood Mac | 235.25 | 7 |
| Ruby | Kaiser Chiefs | 206.08 | 13 |
| Wake Me Up When September Ends | Green Day | 287.14 | 9 |
| Shiver | Coldplay | 302.28 | 12 |
| Summer of ‘69 | Bryan Adams | 234.72 | 10 |
| **Classical** | | | |
| Symphonie Fantastique, II) A Ball | Berlioz | 192.38 | 14 |
| Blue Danube Waltz | Johann Strauss II | 265.85 | 10 |
| Carl Goes Up | Michael Giacchino | 211.00 | 11 |
| Claire de Lune | Debussy | 266.50 | 11 |
| Concerto No 23 | Mozart | 268.79 | 12 |
| Four Seasons, Spring | Vivaldi | 213.58 | 10 |
| Lux Aeterna | Clint Mansell | 243.87 | 11 |
| Nachtmusik | Mozart | 189.94 | 14 |
| Romeo & Juliet | Tchaikovsky | 209.91 | 10 |
| Swan Lake Waltz | Tchaikovsky | 221.42 | 8 |
| **Vocals** (extracted from the rock songs, silences reduced in duration manually) | | | |
| Mr. Brightside | The Killers | 195.50 | 11 |
| Call Me | Blondie | 177.22 | 9 |
| Kids in America | The Muffs | 147.93 | 13 |
| Long Road to Ruin | Foo Fighters | 189.62 | 13 |
| Livin’ On A Prayer | Bon Jovi | 210.16 | 9 |
| Go Your Own Way | Fleetwood Mac | 235.25 | 8 |
| Ruby | Kaiser Chiefs | 170.90 | 13 |
| Wake Me Up When September Ends | Green Day | 200.75 | 14 |
| Shiver | Coldplay | 268.64 | 8 |
| Summer of ‘69 | Bryan Adams | 197.03 | 13 |
| **Speech** (From an audiobook of *The Old Man and the Sea* by Ernest Hemingway, segmented into 7 trials so that trials start and end at silences. The segmentation of the | | | |

| | | | |
|---|---|---|---|
| trials here is identical to the first 7 trials of the Natural Speech dataset: Broderick et al, 2019). | | | |
| audio1 | | 178.06 | 16 |
| audio2 | | 181.20 | 16 |
| audio3 | | 180.57 | 16 |
| audio4 | | 181.85 | 16 |
| audio5 | | 180.83 | 16 |
| audio6 | | 202.63 | 16 |
| audio7 | | 166.22 | 14 |
